# CRISPR/Cas9 gene editing to make conditional mutants of the human malaria parasite *Plasmodium falciparum*

**DOI:** 10.1101/323360

**Authors:** Heather M. Kudyba, David W. Cobb, Anat Florentin, Michelle Krakowiak, Vasant Muralidharan

## Abstract

Malaria is a significant cause of morbidity and mortality worldwide. This disease, which primarily affects those living in tropical and subtropical regions, is caused by infection with *Plasmodium* parasites. The development of better drugs to combat malaria can be accelerated by improving our understanding of the biology of this complex parasite. Genetic manipulation of these parasites is key to understanding their biology, but historically, the genome of *P. falciparum* has been difficult to manipulate. Recently, CRISPR/Cas9 genome editing has been utilized in malaria parasites, allowing for easier protein tagging, generation of conditional protein knockdowns, and deletion of genes. CRISPR/Cas9 genome editing has proven to be a powerful tool for advancing the field of malaria research. Here, we describe a CRISPR/Cas9 method for generating *glmS*-based conditional knockdown mutants in *P. falciparum*. The method is highly adaptable to other types of genetic manipulations, including protein tagging and gene knockouts.

## INTRODUCTION

Malaria is a devastating disease caused by protozoan parasites of the genus *Plasmodium. P. falciparum*, the most deadly human malaria parasite, causes approximately 445,000 deaths per year, mostly in children under the age of five^1^. *Plasmodium* has an intricate life cycle involving a mosquito vector and a vertebrate host. Humans become infected when an infected mosquito takes a blood meal. The parasite first invades the liver where they grow, develop, and divide for approximately one week. After this time, the parasites are released in the bloodstream where they undergo asexual replication in red blood cells (RBC). Growth of the parasites within the red blood cells are directly responsible for all of the clinical symptoms associated with malaria^2^.

Until recently, production of transgenic *P. falciparum* was a laborious process, involving several rounds of drug selection that took many months and had a high rate of failure. This time-consuming procedure relies on the generation of random DNA breaks in the region of interest and the endogenous ability of the parasite to mend its genome though homologous repair^3–6^. Recently, Clustered Regularly Interspaced Palindromic Repeat/Cas9 (CRISPR/Cas9) genome editing has been successfully utilized in *P. falciparum*^7,8^. The introduction of this new technology in malaria research has been critical for advancing understanding of the biology of these deadly *Plasmodium* parasites. CRISPR/Cas9 allows for specific targeting of genes through the use of guide RNAs (gRNAs) that are homologous to the gene of interest. The gRNA/Cas9 complex recognizes the gene through the gRNA and Cas9 introduces a double-strand break, forcing the organism to initiate repair mechanisms^9,10^. Because *P. falciparum* lacks the machinery to repair DNA breaks via non-homologous end joining, it utilizes homologous recombination mechanisms and integrates transfected homologous DNA templates to repair the Cas9/gRNA induced double-strand break^11,12^.

Here we present a protocol for the generation of conditional knockdown mutants in *P. falciparum* using CRISPR/Cas9 genome editing. The protocol demonstrates usage of the *glmS* ribozyme to conditionally knockdown protein level of PfHsp70× (PF3D7_0831700), a chaperone exported by *P. falciparum* into the host red blood cell (RBC)^13,14^. The *glmS* ribozyme is activated by treatment with glucosamine (which is converted to glucosamine-6-phosphate within cells) to cleave its associated mRNA, leading to reduction in protein^14^. This protocol is easily adapted to utilize other conditional knockdown tools such as destabilization domains or RNA aptamers^4,5,15^. Our protocol details the generation of a repair plasmid consisting of a hemagglutinin (HA) tag and *glmS* ribozyme coding sequence flanked by sequences homologous to the PfHsp70× open reading frame (ORF) and 3′UTR. We also describe the generation of a second plasmid to drive expression of the gRNA. These two plasmids, along with a third plasmid that drives expression of Cas9, are transfected into RBCs and used to modify the genome of *P. falciparum* parasites. Finally, we describe a polymerase chain reaction (PCR)-based technique to verify integration of the tag and *glmS* ribozyme. This protocol is highly adaptable for the modification or complete knockout of any *P. falciparum* genes, enhancing our ability to generate new insights into the biology of the malaria parasite.

## PROTOCOL

**Ethics Statement:** Continuous culture of *P. falciparum* requires the use of human RBCs and we utilize commercially purchased units of blood that are stripped of all identifiers and anonymized. The Institutional Review Board and the Office of Biosafety at the University of Georgia have reviewed our protocols and approved all protocols used in our lab.

## 1. Choose gRNA sequence

1.1. Go to CHOP CHOP (http://chopchop.cbu.uib.no/) and select FASTA target. Under “Target”, paste the 200 base pairs from the 3′ end of the open reading frame (ORF) of a gene and 200 base pairs from the start of the gene′s 3′UTR. Under “In”, select the species to be *P. falciparum* (3D7 v3.0) and select CRISPR/Cas9 under “Using”. Next, click “Find Target Sites”.

1.2. Select a gRNA sequence from the options presented, giving preference to the most efficient gRNA that is closest to your site of modification and that has the fewest off-target sites.

Note: Potential gRNA sequences are identified because they are immediately upstream of a Protospacer Adjacent Motif (PAM), which is required for recruitment of Cas9 to DNA. The sequence that is cloned into pMK-U6, the vector that drives gRNA expression, is the 20 bases immediately upstream of the PAM. The PAM specific for *S. pyogenes* Cas9 is the nucleotide sequence NGG and should not be included in the sequence that is cloned into pMK-U6.

CHOP CHOP visually ranks the gRNA sequences, displaying the best options in green, the less ideal options in amber, and the worst options in red. CHOP CHOP gives each gRNA sequence an efficiency score that is calculated using the most up-to-date parameters found in the literature, and they predict off-target sites that could be recognized by the gRNA. Two or three gRNA sequences may need to be tried to find the gRNA best suited to a particular gene.

1.3.Purchase the gRNA sequence and its reverse-complement as Polyacrylamide Gel Electrophoresis-purified oligos; the gRNA sequence used to target PfHSP70x can be found in **Figure IB.**

**Figure 1:**
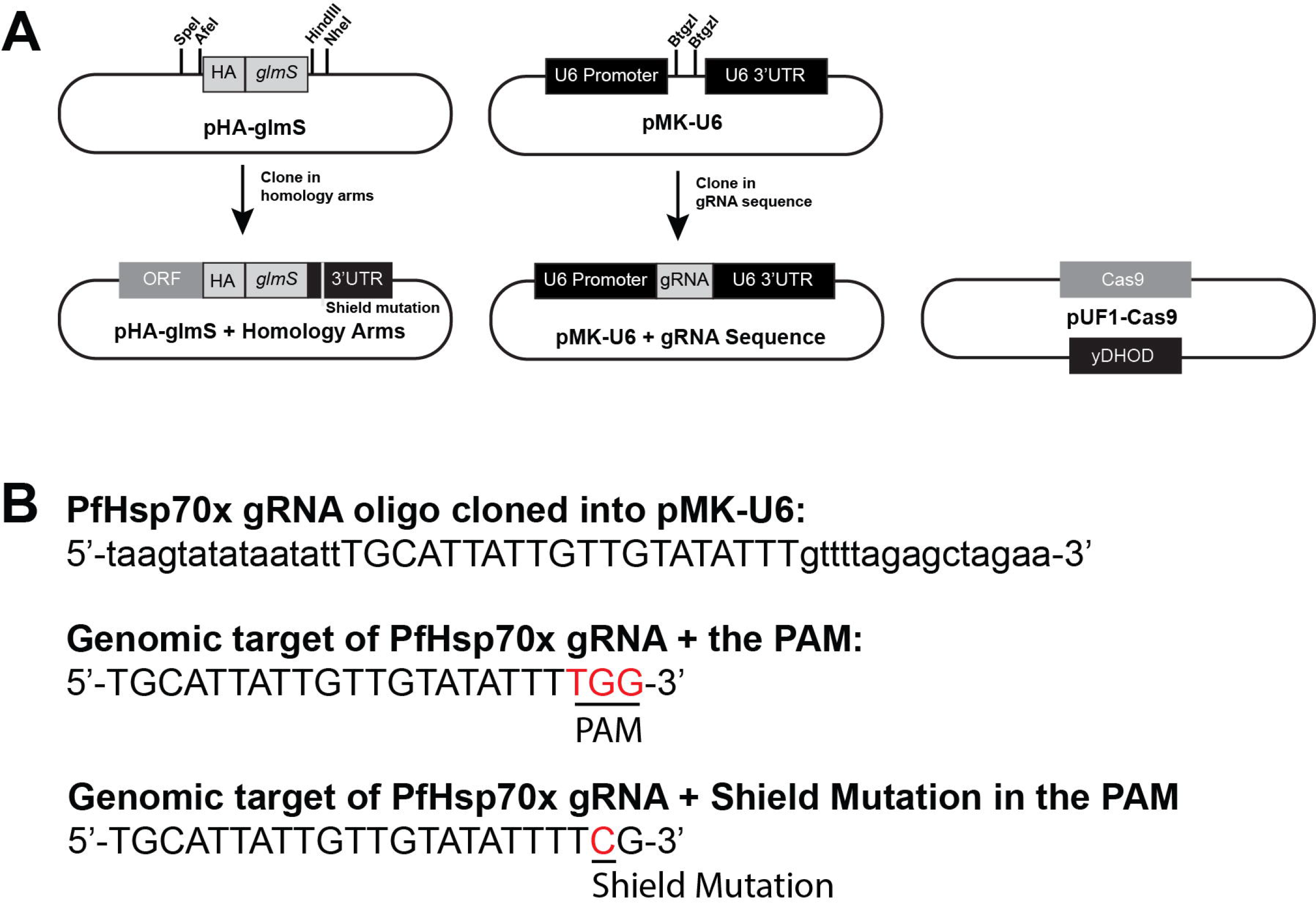
Summary of our three-plasmid approach to CRISPR/Cas9 and examples of a gRNA oligo and shield mutation. A) Schematics of empty pHA-glmS and pMK-U6 are shown with the restriction enzyme sites used for cloning. Also shown are pHA-glmS and pMK-U6 after the homology arms and gRNA sequences have been cloned into them, respectively. Finally, pUF1-Cas9 is shown. yDHOD: yeast dihydrofolate reductase, the resistance marker to DSM1. B) The forward oligo used for cloning the PfHsp70× gRNA sequence into pMK-U6 is shown, with the gRNA sequence in capital letters and the pMK-U6 homology arms necessary for cloning shown in lower case (Top). The genomic target of the PfHsp70× gRNA is shown as well as the downstream PAM, in red (Middle). The shield mutation in the PfHsp70× gRNA PAM is shown in red (Bottom).

Note: This oligo should include 15 base pairs homologous to the gRNA-expressing plasmid, which are necessary for sequence and ligation-independent cloning (SLIC) into the pMK-U6 vector^16^.

## 2. Clone gRNA sequence into pMK-U6

2.1. Digest pMK-U6 with BtgZI.

2.1.1. Digest 10 μg of pMK-U6 with 5 μL of BtgZI enzyme (5000 units/mL) for 3 h at 60 °C. Follow the enzyme manufacturer′s protocol for reaction conditions.

2.1.2. After the 3 h incubation, add an additional 3 μL BtgZI to the reaction to ensure complete digestion of the plasmid. Digest for an additional 3 h, again following manufacturer′s instructions for ensuring the correct reaction condition.

2.1.3. To purify the digested pMK-U6 from the reaction, use a column-based PCR cleanup kit according to manufacturer′s instructions.

2.1.4. Separate the digested DNA using a 0.7% agarose gel and extract the 4,200 base pair band.

2.2. Anneal the oligos containing the gRNA sequence.

2.2.1. Reconstitute the PAGE-purified oligos to a concentration of 100 μM using nuclease free water.

2.2.2. Combine 10 μL of each oligo with 2.2 μL 10× buffer 2 (see the Table of Materials); the total reaction volume will be 22.2 μL.

2.2.3. Run the gRNA annealing program in a thermocycler: Step 1− 95 °C, 10 min; step 2− 95 °C, 1 s, with a reduction in temperature of 0.6 °C/cycle; step 3− Go to step 2,16 times; step 4− 85 °C, 1 min; step 5− 85 °C, 1 s, with a reduction in temperature of 0.6°C/cycle; step 6− Go to step 5,16 times; step 7− 75 °C, 1 min; step 8− 75 °C, 1 s, with a reduction in temperature of 0.6 °C/cycle; step 9− Go to step 8,16 times. Steps 10-21− Repeat the procedure used in Steps 4-9 until the temperature reaches 25 °C; step 22− 25 °C, 1 min.

2.3. Insert the annealed gRNA oligos into the BtgZI-digested and gel-purified pMK-U6 plasmid.

2.3.1. Combine 100 ng digested pMK-U6 with 1 μL 10× buffer 2.1 and 3 μL annealed gRNA oligos. Bring the volume up to 9.5 μL with nuclease-free water.

2.3.2. Add 0.5 μL T4 polymerase and incubate reaction at room temperature for 2 min 30 s.

2.3.3. Move the reaction to ice and incubate for 10 min.

2.3.4. Immediately transform 5 μLofthe reaction into competent *E. coli* according to manufacturer′s instructions and plate bacteria on Lysogeny Broth (LB) agar plates containing 100 μg/mL Ampicillin.

2.3.5. Allow transformed bacteria to grow at 37 °C overnight, then select colonies for DNA extraction with a commercially available plasmid miniprep kit.

## 3. Design homology regions of the repair template

3.1. Design shield mutations within the homology repair template to prevent re-cutting of DNA that is integrated into the genome.

Note: The shield mutation typically consists of introducing a silent mutation to alter the protospacer adjacent motif (PAM) so that Cas9 will not induce a break in the repair template. The PAM required for the Cas9 used here is the nucleotide sequence NGG, where N is any nucleotide. If possible, change one of the G nucleotides to an A, C, or T.

3.1.1. If the PAM cannot be silently mutated, introduce at least two silent mutations into the six base pairs directly adjacent to the PAM^7,8^.

Note: These mutations will prevent recognition of the repair template by the gRNA and prevent re-cutting of the repaired locus by the Cas9/gRNA complex. The shield mutations can be introduced into the homology region by amplifying the DNA with primers that contain the mutation.

3.2. Amplify the ORF homology region for the repair template.

3.2.1. Using PCR, amplify 800 base pairs from the 3′ end of the target gene′s ORF. Design the primers to be used to exclude the stop codon from this amplicon.

3.2.2. Additionally, design the primers to insert this amplicon into pHA*-glmS* that has been digested with Sacll and Afel, either through a DNA ligation reaction or SLIC^16^

3.3. Amplify the 3′UTR homology region for the repair template.

3.3.1. Using PCR, amplify the 800 base pairs immediately following the stop codon of the target gene. The primers used should be designed to insert this amplicon into pHA*-glmS* that has been digested with Hindlll and Nhel, either through a DNA ligation reaction or SLIC^16^.

Note: The high AT content of the *P. falciparum* genome often makes amplification of regions such as UTRs difficult. An alternative approach to using PCR is to synthesize the homology regions.

## 4. Clone homology regions into the repair plasmid

4.1. Insert ORF homology region into pHA *-glmS*.

4.1.1. Digest pHA*-glmS* with Sacll and Afel, according to enzyme manufacturer′s instructions, and insert the ORF homology region PCR product into the digested plasmid using SLIC ^16^.

4.1.2. Transform into competent *E. coli*.

4.2. Insert 3′UTR homology region into pHA-glmS plasmid that already contains the ORF homology region (see 4.1).

4.2.1. Digest the plasmid with Hindlll and Nhel according to enzyme manufacturer’s instructions, and then insert the 3′UTR homology region amplicon into the digested plasmid using SLIC^16^.

4.2.2. Transform into competent *E. coli*. See steps 2.3.4 and 2.3.5 above.

## 5. Precipitate DNA for transfection

5.1. Add 40 μg each of pMK-U6, pUF1-Cas9, and pHA*-glmS* DNA (for a total of 120 μg of DNA) into a sterile 1.5 mL microcentrifuge tube.

5.2. Add 1/10th the volume of DNA of 3M sodium acetate in water (pH 5.2) to the tube and mix well using a vortex. For example, if the volume in step 5.1 was 100 μL, add 10 μL sodium acetate.

5.3. Add 2.5 volumes of 100% ethanol to the tube and mix well using a vortex for at least 30 s. For example, if the volume in 5.1 was 100 μL, add 250 μL 100% ethanol.

5.4. Place the tube on ice or at −20 °C for 30 min.

5.5. Centrifuge the tube at 18,300g for 30 min at 4 °C.

5.6. Carefully remove the supernatant from the tube. Do not disturb the pellet.

5.7. Add 3 volumes of 70% ethanol to the tube and mix briefly using a vortex. For example, if the volume in 5.1 was 100 μL, add 300 μL 70% ethanol.

5.8. Centrifuge the tube at 18,300g for 30 min at 4 °C.

Note: This step should be performed under sterile conditions in a biological safety cabinet.

5.9. Carefully remove the supernatant from the tube. Do not disturb the pellet. Leave the tube open and allow the pellet to air dry for 15 min.

5.10. Store the precipitated DNA at −20 °C until it is needed for transfection.

## 6. Isolate human RBCs from whole blood in preparation for transfection.

6.1. Aliquot fresh blood into sterile 50 mL conical tubes (approximately 25 mL per tube).

6.2. Centrifuge tubes at 1088g for 12 min, with centrifuge brakes set to 4.

6.3. Aspirate off supernatant and buffy coat.

6.4. Resuspend RBC pellet with equal volume incomplete RPMI.

Note: Incomplete RPMI is prepared by supplementing RPMI 1640 with 10.32 μM thymidine, 110.2 μM hypoxanthine, 1 mM sodium pyruvate, 30 mM sodium bicarbonate, 5 mM HEPES,
11.1 mM glucose, and 0.02% (v/v) gentamicin.

6.5. Repeat steps 6.2-6.4 twice.

6.6. After the last wash, resuspend RBCs in equal volume incomplete RPMI and store at 4 °C.

## 7. Transfect RBCs with the CRISPR/Cas9 plasmids (to be done aseptically)

Note: Plasmodium falciparum cultures are maintained as described^17^. Whenever blood is used in the protocol, it is referring to pure red blood cells prepared in Step 6. Blood used should not be older than 6 weeks, as we see a decrease in parasite proliferation in older blood. We describe here a protocol for pre-loading RBCs with DNA and then adding parasite culture to the transfected cells. Other established transfection protocols would be compatible with transfecting these constructs^18,19^.

7.1. Prepare l× cytomix buffer in water (120 mM KCl, 0.15 mM CaCl_2_, 2mM EGTA, 5mM MgCl_2_, 10mM K_2_HPO_4_, 25mM HEPES, pH 7.6). Filter-sterilize the buffer using a 0.22 μM filter.

7.2. Add 380 μL of 1× cytomix to the DNA precipitated in step 5 and vortex to dissolve. Allow the DNA to dissolve in 1× cytomix for 10 minutes, vortexing every 3 minutes for 10 seconds.

7.3. In a sterile 15 mL conical tube, combine 300 μL of red blood cells (RBCs, 50% hematocrit, from Step 6) in incomplete RPMI with 4 mL of 1× cytomix.

7.4. Centrifuge the RBCs from 7.3 at 870g for 3 min, then remove the supernatant from the RBC pellet.

7.5. Resuspend the RBC pellet with DNA/cytomix mixture from step 6.2 and transfer to a 0.2 cm electroporation cuvette.

7.6. Electroporate the RBCs using the following conditions: 0.32kV, 925 μF, capacitance set to “High Cap”, and resistance set to “infinite”.

7.7. Following electroporation, transfer the contents from the cuvette to a 15 mL conical containing 5 mL of complete RPMI (cRPMI). Centrifuge the tube at 870g for 3 min at 20 °C, then decant the supernatant.

Note: cRPMI is prepared as described for incomplete RPMI in 7.4 with the addition of 0.25% (w/v) lipid-rich bovine serum albumin.

7.8. Resuspend the pellet in 4 mL cRPMI and transfer to a well of a 6-well tissue culture plate. Add 400 μL of a high-schizont culture (7-10% schizont parasitemia is ideal) to the transfected RBCs.

Note: Parasitemia is defined as the percentage of parasite-infected RBCs.

7.9. The next day, wash the culture with 4 mL of cRPMI.

7.9.1. Centrifuge the culture at 870g for 3 min and aspirate the supernatant. Resuspend the culture in 4 mL cRPMI.

7.10. 48 h after step 7.6, wash the culture with 4 mL of cRPMI, then resuspend the culture in cRPMI containing 1 μM DSM1 to select for the Cas9 plasmid.

7.11. Continue washing the cultures each day with cRPMI until parasites are no longer visible by blood smear. After this point, the culture medium should be replaced with fresh cRPMI + 1 μM DSM1 every 48 h.

7.11.1. To make a blood smear, pipette 150 μL of culture into an 0.6 mL eppendorf tube. Pellet the cells by centrifugation at 1700g for 30 s.

7.11.2. Aspirate off the supernatant. Use a pipette to transfer the pelleted cells to a glass slide. Using a second glass slide, held at a 45° angle to the first slide, smear the blood droplet. Stain the slide using a commercially available staining kit according to the manufacturer′s protocol.

7.11.3 View parasites using a 100× oil immersion objective.

7.12. Beginning 5 days post-transfection (Step 7.6), remove 2 mL of the culture, with RBCs resuspened in the culture medium, and add back 2 mL fresh medium (cRPMI + 1 μM DSM1) and blood at 2% hematocrit. Add fresh blood in this manner once a week until parasites reappear, as determined by thin blood smear.

Note: If integration is successful, parasites generally reappear in culture by one-month posttransfection.

7.13. Once parasites reemerge, remove drug pressure. Alternatively, remove drug pressure after parasites have been cloned out.

## 8. Check parasites for integration of the repair template

8.1. When parasites are visible again by thin blood smear, isolate DNA from the culture.

8.2. Use PCR to amplify the modified region of the genome to determine whether the targeted locus has been successfully altered and whether the unmodified *wild-type* locus (indicative of *wild-type* parasites) is detectable.

8.2.1. To detect parasites that have integrated the repair template, use a forward primer that sits at the beginning of the ORF, outside of the cloned homology region. Use a reverse primer that sits in the 3′UTR.

Note: As this amplification includes the sequences of the HA tags and *glmS* ribozyme, amplicons from integrated parasites will be longer than the same region amplified in *wild-type* parasites.

## 9. Clone parasites by limiting dilution

9.1 Perform serial dilutions of the parasite culture from step 7.13 to achieve a final concentration of 0.5 parasites/200 μL.

9.1.1. Prepare 1 mL culture in cRPMI at 5% parasitemia and 2% hematocrit; at this parasitemia and hematocrit, the culture contains 1×10^7^ parasites/mL.

9.1.2. Dilute this culture 1:100 with cRPMI. Dilute again 1:100 with cRPMI.

9.1.3. Dilute 1:400. Perform this dilution by adding 62.5 μL culture to 25 mL cRPMI and 1 mL blood. This dilution results in the desired concentration of 0.5 parasites/200 μL.

9.1.5. Add 200 μL of the diluted culture to the wells of a 96-well tissue culture plate.

9.2. Maintain the cloning plate until parasites are detectable in the wells.

9.2.1. Every 48 h, replace the medium in the 96-well plate with fresh medium.

9.2.2. Once a week, starting 5 days after beginning the cloning plate (Step 8.1.5), remove 100 μL from each well and add back 100 μL of fresh medium + blood (2% hematocrit).

9.3. Identify wells containing parasites.

9.3.1. Place the 96-well plate at a 45° angle for approximately 20 min, allowing the blood to settle at an angle within the plate.

9.3.2. Place the 96-well plate on a light box. Observe that the wells containing parasites will contain medium that is yellow in color, compared to the pink medium of parasite-free wells, due to acidification of the medium by the parasites.

9.3.3. Using a serological pipette, move the contents of the parasite-containing wells to a 24− well tissue culture plate to allow expansion of parasitemia.

9.3.4. Using PCR analysis as described in Step 8, check these clonal parasite lines for correct integration.

## 10. Knockdown protein by treating parasites with GlcN and confirm knockdown by Western blot analysis

10.1. Prepare 0.5 M GlcN stock solution; the stock can be stored at −20° C.

10.2. Add GlcN to *glmS* parasite cultures and allow to grow in the presence of GlcN.

Note: The final concentration and timing of GlcN treatment to be used is dependent upon the experiment and parasite line. GlcN can impact parasite growth, so the parental parasite strain should be exposed to a range of GlcN concentrations to determine sensitivity to the compound. Often, a concentration of 2.0-7.5 mM GlcN is used^13,14,20^.

10.3. Isolate protein samples from GlcN-treated parasites^13^.

10.4. Use protein samples for Western blot analysis to detect reduction in protein^13^.

10.4.1. Use an anti-HA antibody according to manufacturer′s instructions to detect the HA-glmS-tagged protein, and compare the HA band to a loading control, such as PfEFla.

## REPRESENTATIVE RESULTS

A schematic of the plasmids used in this method as well as an example of a shield mutation are shown in **Figure 1.** As an example of how to identify mutant parasites after transfection, results from PCRs to check integration of the HA *-glmS* construct is shown in **Figure 2.** A representative image of a cloning plate is shown in **Figure 3** to demonstrate the color change of the medium in the presence of parasites. Results from an immunofluorescence assay and Western blotting experiments are shown in **Figure 4** to demonstrate the functionality of the HA tag and *glmS-* based reduction of protein in the parasites. **Figure 5** demonstrates the inability of short homology arms on PCR products to modify the parasites genome and obtain viable mutants.

**Figure 2:**
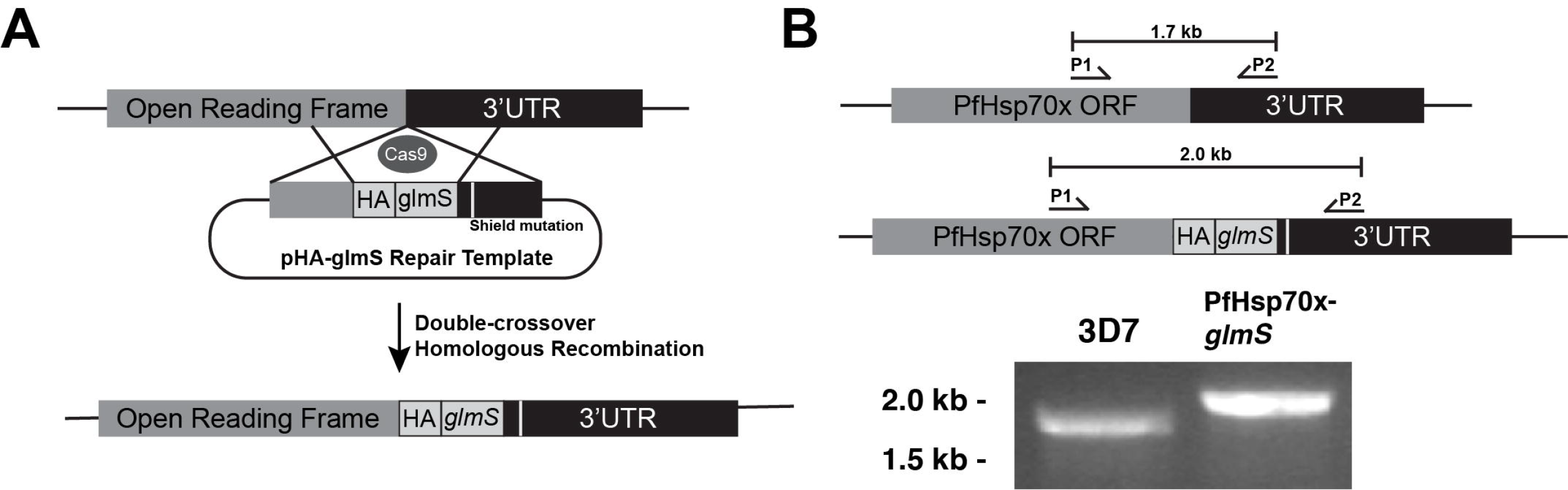
Schematic of CRISPR/Cas9 genome modification using pHA-glmS and strategy for confirming integration. A) Cas9, guided to a genomic locus by a gRNA, induces a double strand break in the DNA. The parasite repairs the damage through double crossover homologous repair, using the pHA-glmS plasmid as a template and introducing the HA-glmS sequence into the genome. B) A PCR test to identify correct integration of the HA-glmS sequence. Using primers P1 and P2, the 3′ ORF of *wild-type* PfHsp70× and PfHsp70×-g/mS mutants are amplified^13^. The amplicon from PfHsp70×-glmS is longer than *wild-type* due to insertion of the HA-glmS sequence.

**Figure 3:**
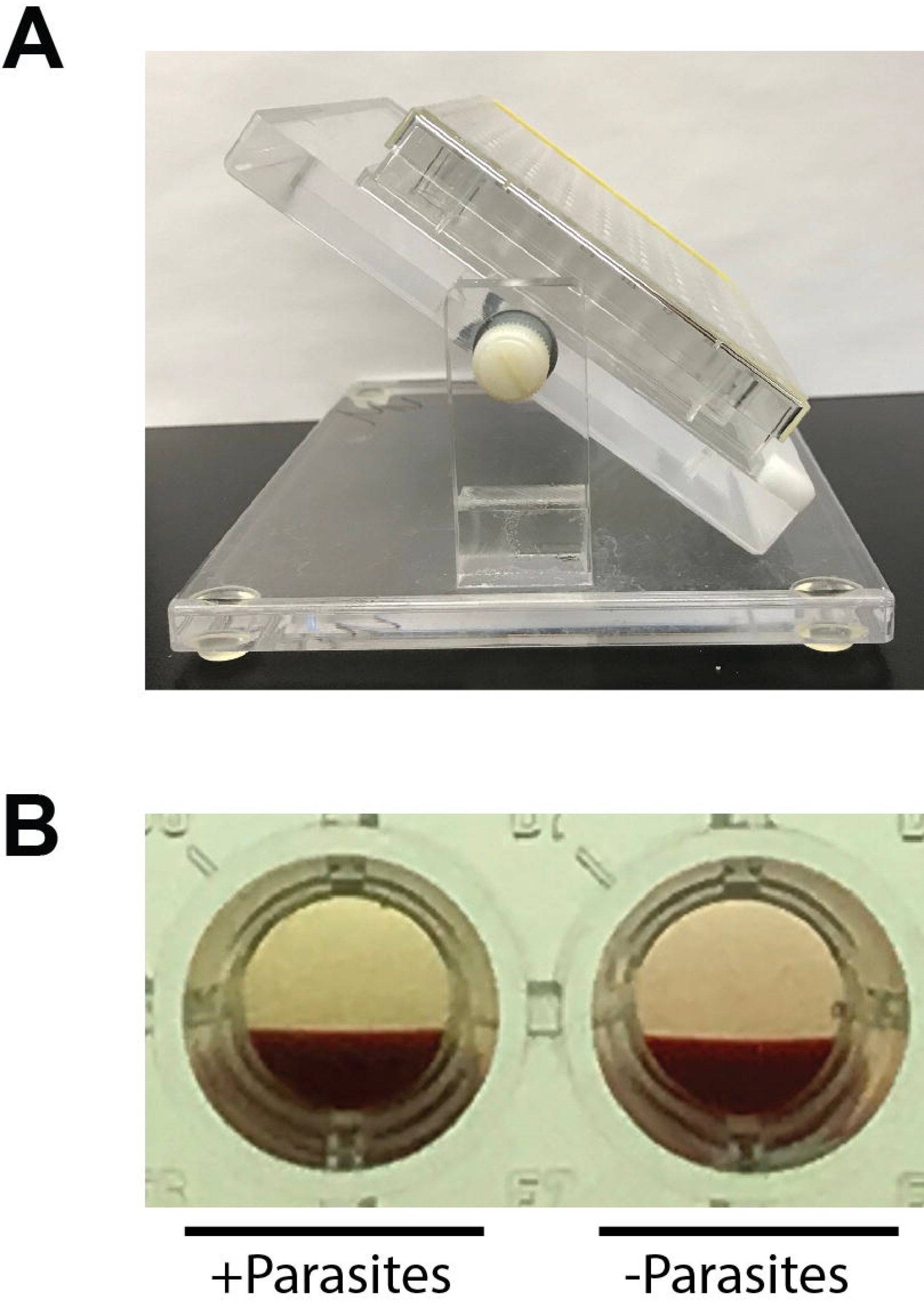
Identification of wells containing parasites in a 96-well cloning plate. A) The 96-well plate is set at a 45° angle for approximately 20 minutes to allow the blood to settle at an angle in the plate. B) The well on the left contains a parasite culture, indicated by the yellow color of the medium in comparison to the pink medium of the parasite-free well on the right.

**Figure 4:**
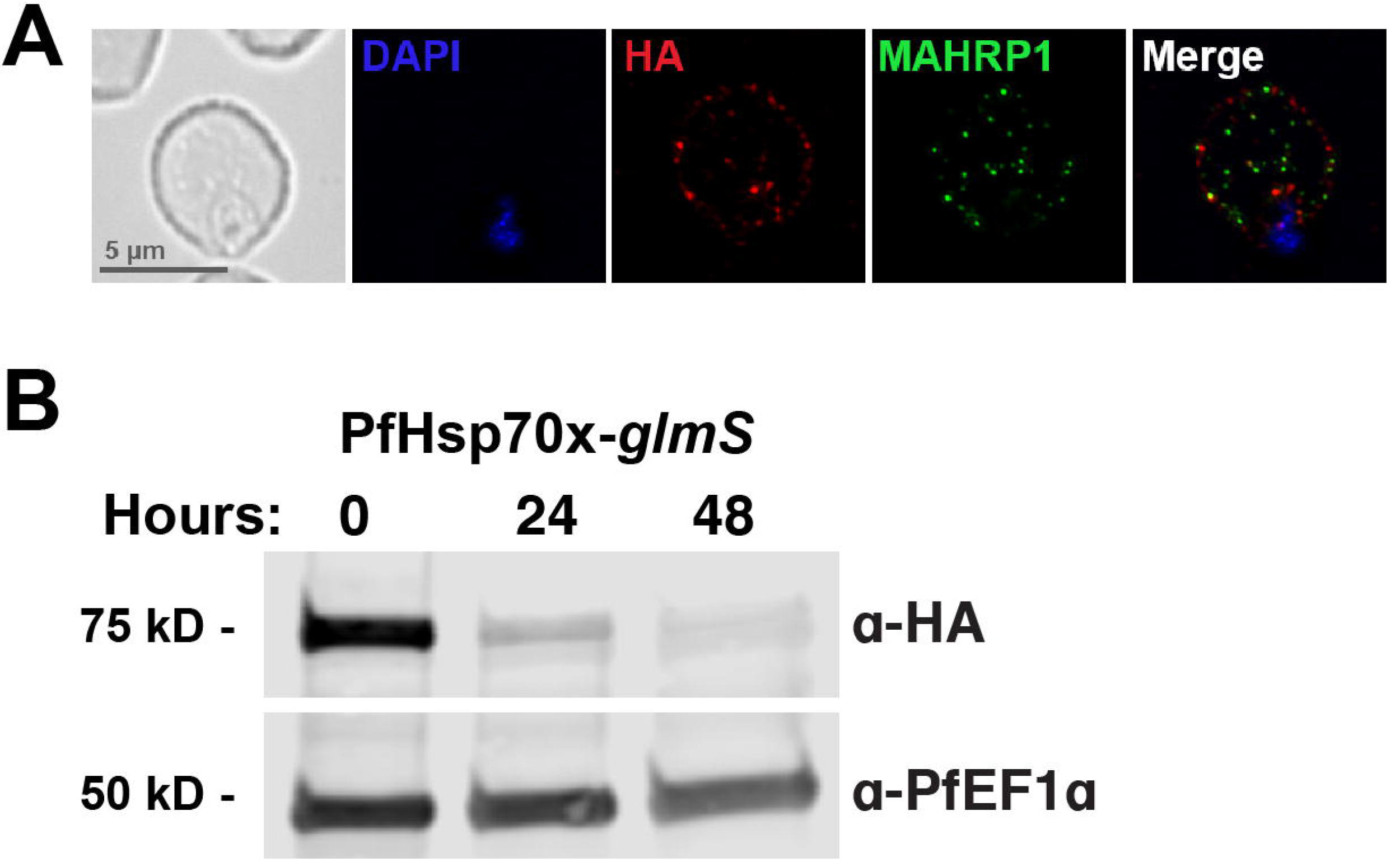
An immunofluorescence assay shows the correct HA-tagging of PfHsp70× and Western blotting shows reduction of PfHsp70x protein levels during treatment with glucosamine. A) PfHsp70×-*glmS* parasites were fixed and stained with DAPI (nucleus marker) and antibodies to HA and MAHRP1 (Membrane Associated Histidine Rich Protein 1, a marker of protein export to the host RBC)^13^.B) PfHsp70×-*glmS* parasites were treated with 7.5 mM glucosamine and whole-parasite lysates were used for Western blotting analysis^13^. The membrane was probed with antibodies for HA and PfEF1α as a loading control^13^. As expected, glucosamine treatment results in a reduction of the protein.

**Figure 5:**
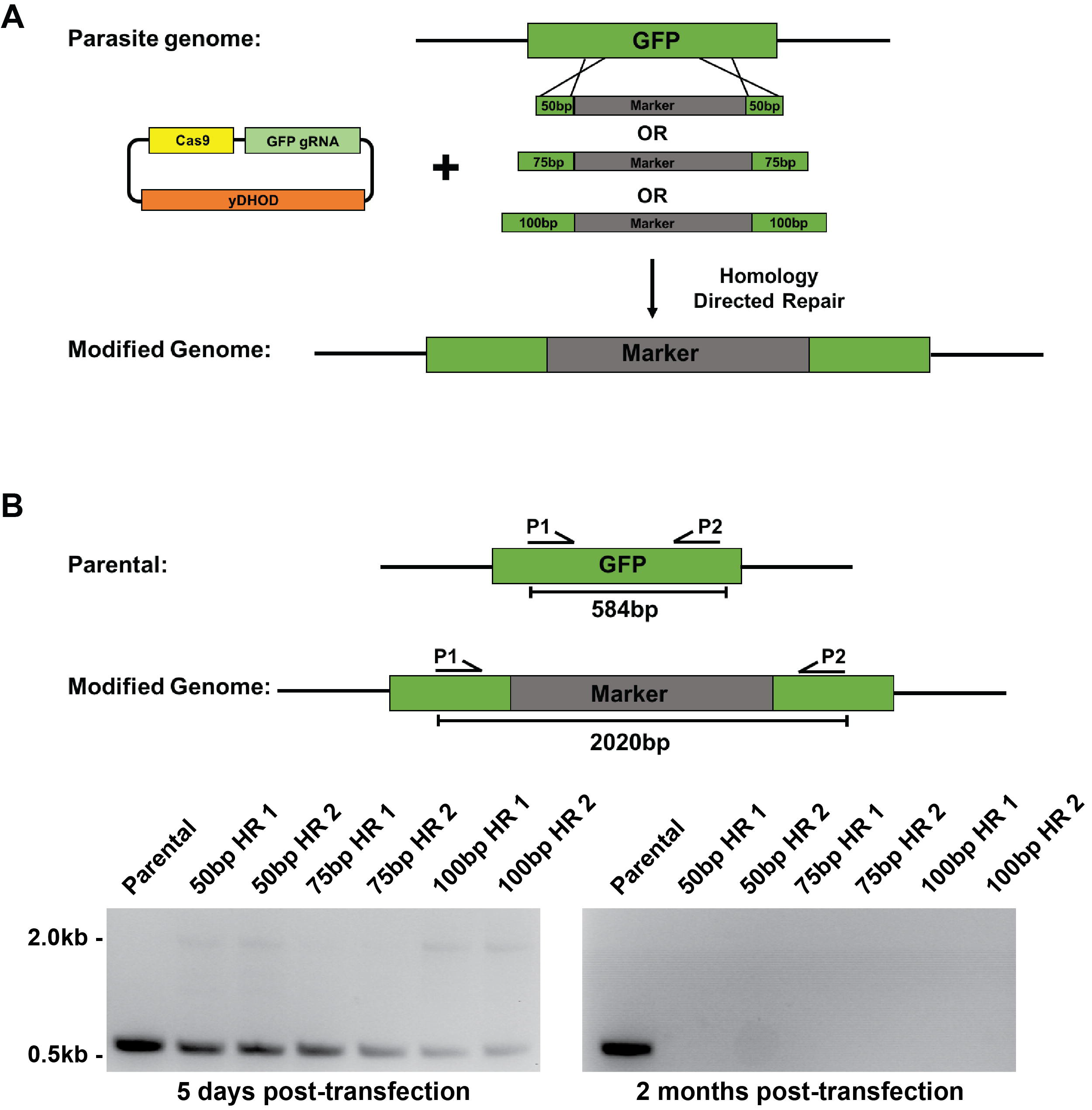
Using short homology sequences for repair. A) Schematic representation showing knockout of GFP in B7 parasites^21^. B7 parasites are a derivative of 3D7 where Plasmepsin II has been tagged with GFP. PCR products containing 50, 75, or 100 base pairs of GFP homology regions flanking a blasticidin S resistance cassette (labeled “marker”), along with pUF1-Cas9-eGFP-gRNA, a plasmid expressing Cas9 and a GFP gRNA, were transfected into B7 parasites. Each transfection was carried out twice. Drug pressure (DSM1) was applied 2-day post transfection. B) PCR test on DNA isolated from transfected parasites five days post-transfection and two months post-transfection. Primers used to test integration of the BSD resistance cassette will yield a 584 base pair product for B7 parental parasites and a 2020 base pair product for parasites that have integrated the marker.

## DISCUSSION

The implementation of CRISPR/Cas9 in *P. falciparum* has both increased the efficiency of and decreased the amount of time needed for modifying the parasite’s genome, in comparison to previous methods of genetic manipulation. The comprehensive protocol described in this manuscript outlines the steps taken to generate conditional mutants using CRISPR/Cas9 in *Plasmodium falciparum*. While the method here is written specifically for the generation of HA-*glmS* mutants, this strategy can be adapted for a variety of purposes, including the tagging of genes, gene knockouts, and introduction of point mutations.

A critical early step in this protocol is the selection of a gRNA sequence. When selecting a gRNA, there are several considerations to keep in mind in regards to where the gRNA sits, how efficient it is, and whether it has the potential for off-target effects. Typically, the gRNA sequence should be as close as possible to the site of modification, ideally within 200 bp. This will decrease the likelihood of the parasites using the repair template to fix their genome without integrating the tag. The tool used here to locate a gRNA was a free online service called CHOP CHOP^22^. Another online tool, Eukaryotic Pathogen CRISPR guide RNA/DNA Design Tool (EuPaGDT, http://grna.ctegd.uga.edu/), can also be used^23^. EuPaGDT provides additional characterization of gRNA sequences, including prediction of off-target hits and potential issues that may prevent transcription of the gRNA. EuPaGDT also has tools for batch processing of gRNAs to target multiple genes or whole genomes. The gRNA chosen should be one that sits closest to the site of modification with the highest efficiency and minimal off-target hits. An important limitation to CRISPR/Cas9 gene editing that may arise is the inability to design a suitable gRNA to target the gene of interest. In such cases, a trial-and-error approach may be needed, using multiple sub-optimal gRNA sequences until the best one is found and successful gene editing has occurred.

Another important consideration for generating *P. falciparum* mutants using CRISPR/Cas9 is the length of the homology regions used in the repair template. The protocol here states that the homology regions should be approximately 800 base pairs each, but we have also been successful in using smaller regions (500 base pairs)^3^. Successful genome modification using CRISPR/Cas9 and short homology arms on PCR products have been used in other protozoan parasites such as *Toxoplasma gondii* and *Trichomonas vaginalis*^24,25^. We tested the feasibility of using smaller homology arms on PCR products (50, 75, or 100 base pairs) by attempting to knockout GFP in B7 parasites using a blasticidin resistance cassette^21^. We saw some integration of the blasticidin resistance cassette at five days post transfection; however, these parasites never recovered from transfection. For these transfections, we selected for the Cas9-expressing plasmid using DSM1. A different selection method, such as treating transfected cultures with blasticidin S alone or in combination with DSM1, may improve the chances of parasites reappearing when using shorter homology regions for repairing the Cas9/gRNA induced break. We did not select with blasticidin S in this case because we wanted to test whether short homology arms could be used in instances where a drug resistance cassette is not being integrated into the genome, such as when a protein is being tagged.

The core components of CRISPR/Cas9 gene editing discussed here are the Cas9 endonuclease, the gRNA, and the repair template. We describe a three-plasmid approach to introduce these components into the parasites, where Cas9, the gRNA, and the repair template are found in separate plasmids. In addition to this approach, our lab has been successful in using a two-plasmid approach where Cas9 and gRNA expression are driven by a single plasmid and the repair template is found in a second plasmid^29^. Furthermore, a few labs are using a strain of *Plasmodium* (NF54^attB^) which constitutively expresses Cas9 and a T7 RNA polymerase to drive expression of gRNA′s^30^. In this case, a single plasmid containing the repair template and the gRNA are transfected into NF54^attB^ parasites^31,32^. A plasmid-free approach, utilizing a purified Cas9-gRNA ribonucleoprotein complex, has been used to insert mutations into the genome as well^33^. The success of these different approaches demonstrates flexibility in how researchers can introduce the Cas9/gRNA components into the parasite.

Finally, the choice of drug pressure to apply to transfected parasites can be altered depending on constructs used. Here, we show successful generation of mutants by transiently selecting for the Cas9 expressing plasmid using DSM1 until parasites reappear. To generate PfHsp70x knockout parasites, *pfhsp70×* was replaced with the human dihydrofolate reductase gene, and parasites were selected using WR99210^13^.The recently described TetR-PfDOZI knockdown system relies on integration of a plasmid containing a blasticidin S resistance gene, allowing for selection of parasites using blasticidin S^15,31^.

CRISPR/Cas9 gene editing of *P. falciparum* has proven to be a powerful tool in malaria research, and we have detailed here a method for generating conditional knockdown mutants^3,7,8,13,20,28^. The protocol is highly adaptable to individual research interests.

## ACKNOWLEDGMENTS

We thank Muthugapatti Kandasamy at the University of Georgia Biomedical Microscopy Core for technical assistance and Jose-Juan Lopez-Rubio for sharing the pUF1-Cas9 and pL6 plasmids. This work was supported by ARCS Foundation awards to D.W.C. and to H.M.K., UGA startup funds to V.M., grants from the March of Dimes Foundation (Basil O′Connor Starter Scholar Research Award) to V.M., and the US National Institutes of Health (R00AI099156 and R01AI130139) to V.M. and (T32AI060546) to H.M.K.

## REFERENCES

1 World Health Organization. World Malaria Report. WHO, Geneva. (2017).

2 Miller, L. H., Baruch, D. I., Marsh, K. & Doumbo, O. K. The pathogenic basis of malaria. Nature. 415 (6872), 673–679, (2002).

3 Florentin, A. et al. PfClpC Is an Essential Clp Chaperone Required for Plastid Integrity and Clp Protease Stability in Plasmodium falciparum. Cell Rep. 21 (7), 1746–1756, (2017).

4 Muralidharan, V., Oksman, A., Pal, P., Lindquist, S. & Goldberg, D. E. Plasmodium falciparum heat shock protein 110 stabilizes the asparagine repeat-rich parasite proteome during malarial fevers. Nat Commun. 3 1310–1310, (2012).

5 Muralidharan, V., Oksman, A., Iwamoto, M., Wandless, T. J. & Goldberg, D. E. Asparagine repeat function in a Plasmodium falciparum protein assessed via a regulatable fluorescent affinity tag. Proc Natl Acad Sci U S A. 108 (11), 4411–4416, (2011).

6 Beck, J. R., Muralidharan, V., Oksman, A. & Goldberg, D. E. PTEX component HSP101 mediates export of diverse malaria effectors into host erythrocytes. Nature. 511 (7511), 592–595, (2014).

7 Ghorbal, M. et al. Genome editing in the human malaria parasite Plasmodium falciparum using the CRISPR-Cas9 system. Nat Biotechnol. 32 (8), 819–821, (2014).

8 Wagner, J. C., Platt, R. J., Goldfless, S. J., Zhang, F. & Niles, J. C. Efficient CRISPR-Cas9-mediated genome editing in Plasmodium falciparum. Nat Methods. 11(9), 915–918, (2014).

9 Doudna, J. A. & Charpentier, E. Genome editing. The new frontier of genome engineering with CRISPR-Cas9. Science. 346 (6213), 1258096, (2014).

10 Wang, H., La Russa, M. & Qi, L. S. CRISPR/Cas9 in Genome Editing and Beyond. Annu Rev Biochem. 85 227–264, (2016).

11 Kirkman, L. A. & Deitsch, K. W. Antigenic variation and the generation of diversity in malaria parasites. Curr Opin Microbiol. 15 (4), 456–462, (2012).

12 Lee, A. H., Symington, L. S. & Fidock, D. A. DNA Repair Mechanisms and Their Biological Roles in the Malaria Parasite Plasmodium falciparum. Microbiology and Molecular Biology Reviews. 78 (3), 469–486, (2014).

13 Cobb, D. W. et al. The Exported Chaperone PfHsp70× Is Dispensable for the Plasmodium falciparum Intraerythrocytic Life Cycle. mSphere. 2 (5), (2017).

14 Prommana, P. et al. Inducible knockdown of Plasmodium gene expression using the glmS ribozyme. PLoS One. 8 (8), e73783, (2013).

15 Ganesan, S. M., Falla, A., Goldfless, S. J., Nasamu, A. S. & Niles, J. C. Synthetic RNA-protein modules integrated with native translation mechanisms to control gene expression in malaria parasites. Nat Commun. 7 10727, (2016).

16 Li, M. Z. & Elledge, S. J. Harnessing homologous recombination in vitro to generate recombinant DNA via SLIC. Nat Methods. 4 (3), 251–256, (2007).

17 Drew, M. E. et al. Plasmodium food vacuole plasmepsins are activated by falcipains. J Biol Chem. 283 (19), 12870–12876, (2008).

18 Wu, Y., Sifri, C. D., Lei, H.-H., Su, X.-Z. & Wellems, T. E. Transfection of Plasmodium falciparum within human red blood cells. Proc Natl Acad Sci U S A. 92 973–977, (1995).

19 Janse, C. J. et al. High efficiency transfection of Plasmodium berghei facilitates novel selection procedures. Mol Biochem Parasitol. 145 (1), 60–70, (2006).

20 Counihan, N. A. et al. Plasmodium falciparum parasites deploy RhopH2 into the host erythrocyte to obtain nutrients, grow and replicate. Elife. 6, (2017).

21 Klemba, M., Beatty, W., Gluzman, I. & Goldberg, D. E. Trafficking of plasmepsin II to the food vacuole of the malaria parasite Plasmodium falciparum. J Cell Biol. 164 (1), 47–56, (2004).

22 Labun, K., Montague, T. G., Gagnon, J. A., Thyme, S. B. & Valen, E. CHOPCHOP v2: a web tool for the next generation of CRISPR genome engineering. Nucleic Acids Res. 44 (W1), W272–276, (2016).

23 Peng, D. & Tarleton, R. EuPaGDT: a web tool tailored to design CRISPR guide RNAs for eukaryotic pathogens. Microb Genom. 1 (4), e000033, (2015).

24 Shen, B., Brown, K. M., Lee, T. D. & Sibley, L. D. Efficient Gene Disruption in Diverse Strains of Toxoplasma gondii Using CRISPR/CAS9. MBio. 5 (3), (2014).

25 Janssen, B. D. et al. CRISPR/Cas9-mediated gene modification and gene knock out in the human-infective parasite Trichomonas vaginalis. Sci Rep. 8 (1), 270, (2018).

26 Spillman, N. J., Beck, J. R., Ganesan, S. M., Niles, J. C. & Goldberg, D. E. The chaperonin TRiC forms an oligomeric complex in the malaria parasite cytosol. Cell Microbiol. 19 (6), (2017).

27 Brancucci, N. M. B. et al. Lysophosphatidylcholine Regulates Sexual Stage Differentiation in the Human Malaria Parasite Plasmodium falciparum. Cell. 10.1016/j.cell.2017.10.020, (2017).

28 Ng, C. L. et al. CRISPR-Cas9-modified pfmdr1 protects Plasmodium falciparum asexual blood stages and gametocytes against a class of piperazine-containing compounds but potentiates artemisinin-based combination therapy partner drugs. Mol Microbiol. 101 (3), 381–393, (2016).

29 Lim, M. Y. et al. UDP-galactose and acetyl-CoA transporters as Plasmodium multidrug resistance genes. Nat Microbiol. 10.1038/nmicrobiol.2016.166 16166, (2016).

30 Adjalley, S. H. et al. Quantitative assessment of Plasmodium falciparum sexual development reveals potent transmission blocking activity by methylene blue. PNAS. 108 (47), E1214–E1223, (2011).

31 Sidik, S. M. et al. A Genome-wide CRISPR Screen in Toxoplasma Identifies Essential Apicomplexan Genes. Cell. 166 (6), 1423–1435 e1412, (2016).

32 Amberg-Johnson, K. et al. Small molecule inhibition of apicomplexan FtsH1 disrupts plastid biogenesis in human pathogens. Elife. 6, (2017).

33 Crawford, E. D. et al. Plasmid-free CRISPR/Cas9 genome editing in Plasmodium falciparum confirms mutations conferring resistance to the dihydroisoquinolone clinical candidate SJ733. PLoS One. 12 (5), e0178163, (2017).

